# CAZyme Annotation of Five Parasitic Trichomonads Reveals a Conserved Repertoire of Host Glycan Targeting Proteins

**DOI:** 10.64898/2026.10.12.758501

**Authors:** Adam J. Hart, David N. Bolam, Manuel Banzhaf, Bernard Henrissat, Robert P. Hirt

## Abstract

Deciphering the functional potential of the unusually large and repetitive genomes of trichomonads has remained challenging since the first *Trichomonas vaginalis* genome assembly in 2007. These genomes are characterised by extensive gene duplication, widespread lateral gene transfer from prokaryotes, and numerous hypothetical proteins. Here, we systematically re-annotated five parasitic trichomonad genomes using the CAZy annotation pipeline, identifying 1,406 CAZyme-encoding genes (which encompass all four core CAZyme classes and carbohydrate-binding modules (CBMs)) - including 516 genes previously annotated as encoding hypothetical proteins - and generating predictions of CAZy genes originating from potential lateral gene transfer, encoding candidate secreted proteins, and their candidate metabolic pathway involvement. Comparative analyses revealed a conserved, highly duplicated repertoire of candidate mucin O-glycan-targeting CAZymes across all five species, alongside broad conservation of a candidate antibacterial CAZyme toolkit. Together, these findings substantially enrich functional annotation of these genomes and highlight in particular CAZymes and CBMs as promising targets for the future characterisation of the molecular and cellular basis of parasite-host and parasite-microbiota interactions underlying mucosal disease and dysbiosis associated with trichomonads.

**Significance Statement:** Parasitic trichomonads possess unusually large and complex genomes that are challenging to annotate, hindering the identification of virulence factors and other genes of interest. Glycans are central to interactions between these parasites and their hosts and host microbiota, influencing processes such as attachment, nutrient acquisition and microbial competition. This study enhances the annotation of five parasitic trichomonad genomes by incorporating CAZy-classified protein and module information (e.g. glycoside hydrolases and carbohydrate binding modules) and comparative analyses, revealing novel candidate virulence factors within this functionally important gene set. The resulting dataset provides a rich and robust resource for the research community to further investigate the biology and the molecular and cellular basis of the pathogenic potential of these understudied parasites that infect various mucosal sites across a broad range of hosts across birds and mammals, including humans.

## Introduction

Carbohydrate-active enzymes (CAZymes) are ubiquitous across cellular life, reflecting the exceptional structural diversity of carbohydrates, which constitute some of the most diverse substrates in nature (Laine, 1994; Varki *et al*., 2022). The CAZy database enables the systematic identification of CAZyme and CBM domains across the tree of life through sequence similarity comparisons to a comprehensive, functionally characterised set of curated entries (Drula *et al*., 2022). CAZyme domains are classified into families according to their general catalytic or functional roles: glycoside hydrolases (GH), glycosyltransferases (GT), polysaccharide lyases (PL), carbohydrate esterases (CE), carbohydrate-binding modules (CBM) and Auxiliary Activities (AA) (Lombard *et al*., 2014; Drula *et al*., 2022). Incorporating CAZyme domain annotations into genome annotations enhances the functional characterisation of annotated genes and assigns putative functions to previously hypothetical proteins. This, in turn, improves our understanding of an organism’s capacity to degrade and modify polysaccharides, processes that often confer metabolic advantages and facilitate adaptation to specific ecological niches (Grondin *et al*., 2017; Wardman *et al*., 2022). Additionally, the CAZyme repertoire of a mucosal microbiota can influence the health of the host, and some microbial CAZymes are themselves considered virulence factors in disease (Lammerts van Bueren *et al*., 2011; Cockburn & Koropatkin, 2016; Carlson *et al*., 2018; Segui-Perez *et al*., 2025). Therefore, characterising the CAZyme repertoire of organisms residing at mucosal surfaces, including members of the resident microbiota and pathogens, can help elucidate mechanisms involved in mucosal colonisation, host–microbe interactions and, in the case of pathogens, virulence (Wardman *et al*., 2022; Raba & Luis, 2023).

Mucosal parasitic trichomonads typically have relatively large genomes with extensive gene duplication, features that have been associated with their parasitic lifestyle, virulence traits, and capacity for host or mucosal site switching (Barratt *et al*., 2016; Abdel-Glil *et al*., 2024; Mpeyako *et al*., 2024; Sullivan *et al*., 2025). Since the publication of the first genome assembly and annotation for a parasitic trichomonad, *Trichomonas vaginalis*, substantial effort has gone into understanding why these genomes are so large and how their expanded gene repertoires contribute to parasitism and disease (Carlton *et al*., 2007; Hirt, 2013; Hirt & Sherrard, 2015; Barratt *et al*., 2016; Hirt, 2019). Additionally, these genomes feature a relatively large number of candidate genes originating through lateral gene transfer from prokaryotes (amongst others, e.g. viruses) with which they share a niche (Carlton *et al*., 2007; Sullivan *et al*., 2025).

### Trichomonas spp

CAZymes in particular have been implicated in virulence and modulation of the host environment in several studies. Genome annotation of *Trichomonas tenax* and comparative analysis with *Trichomonas vaginalis* highlighted that conserved CAZymes may contribute to virulence by targeting host glycans such as mucins (Mpeyako *et al*., 2024). Transcriptomic analysis of the avian parasite *Trichomonas gallinae* in co-culture with *E. coli* highlighted a conserved (amongst *Trichomonas*) and duplicated antibacterial toolkit made up primarily of GH CAZymes, which target the bacterial cell wall (Hart *et al*., 2025). The same study showed, using phylogenetic analyses, that several of these CAZymes are conserved across several species, with homologues specifically identified among the parasites *Trichomonas vaginalis*, *Trichomonas tenax*, *Histomonas meleagridis* and *Tritrichomonas foetus* (Hart *et al*., 2025). These bacterial cell wall-targeting GHs likely act in concert with other conserved peptidoglycan hydrolases that cleave peptide cross-links in the bacterial cell wall (Pinheiro *et al*., 2018; Barnett *et al*., 2023). This toolkit may contribute to the initiation or perpetuation of dysbiotic microbiota, which frequently accompanies *Trichomonas* infection (Brotman *et al*., 2012; Ji *et al*., 2020).

Together, these findings suggest that CAZyme-mediated modification of both host mucins and bacterial cell walls is a key feature of parasitic trichomonads and may play an important role in shaping mucosal dysbiosis during infection (Hart *et al*., 2025). This study, for the first time, specifically reanalyses five parasitic trichomonad genomes which reside in different hosts and host mucosal sites (**Table 1**) to comprehensively identify and functionally annotate their CAZyme and CBM repertoires. The identified CAZymes were then evaluated for their likelihood of origin from lateral gene transfer, predicted secretion (via signal peptide detection), KEGG pathway annotation, and membership in orthologous groups. Collectively, this reannotation added CAZy domain assignments to over 1,400 protein-coding genes, including new functional annotation for 516 previously hypothetical or undefined genes. Notably, this analysis confirms the conservation across all five considered trichomonads of the antibacterial CAZyme toolkit previously described in *Trichomonas gallinae* and the mucin-targeting repertoire of *Trichomonas vaginalis* and *Trichomonas tenax*. Moreover, it has highlighted the expanded host glycan targeting capacity of these parasites with potential implications for dysbiosis and disease. The CAZymes identified here therefore represent promising targets for future functional characterisation to clarify how parasitic trichomonads contribute to pathogenesis at mucosal sites.

**Table 1:** Summary of known trichomonad host species and the mucosal sites they inhabit.

| Species | Host/s | Site/s |
| --- | --- | --- |
| *Trichomonas vaginalis* | Human (Hirt & Sherrard, 2015) | Urogenital |
| *Trichomonas gallinae* | Columbiformes, Passeriformes and Falconiformes (Birds) (Amin *et al.*, 2014) | Oral and Gastrointestinal |
| *Trichomonas tenax* | Humans, Companion animals (Dogs, cats) and Birds (Kellerová & Tachezy, 2017; Matthew *et al.*, 2023) | Oral |
| *Histomonas meleagridis* | Chicken, Turkey, other Gallinaceous Birds and an intermediate host *Heterakis gallinarum* (nematode) (Beer *et al.*, 2022) | Gastrointestinal (in birds) and in eggs of *Heterakis gallinarum* |
| *Tritrichomonas foetus* | Cows, Cats and Pigs (Dąbrowska *et al.*, 2019) | Urogenital (Cows), Gastrointestinal (Cats, Cows and Pigs) |

## Results

### CAZyme Annotation Overview

CAZyme repertoires in the five trichomonad species are dominated by GHs and GTs, with a subset of conserved core domains and species-specific expansions, particularly in *Tritrichomonas foetus*. Across the five trichomonad species analysed for CAZyme content, we identified 1,406 unique protein-coding genes containing one or more CAZy domains. Of these, approximately 62% (n=874) belong to glycoside hydrolase families, including GHs that also carry associated CBMs (**Figure 1A-F**). An additional ∼33% (n=463) are GTs. The remaining annotations include 43 CBMs, 3 CEs, and 23 PLs. Among the 80 distinct CAZy domain families identified, 31 are conserved across all five species (**Figure 1G**). Additionally, for the 1,406 identified CAZymes, 209 also have sub-family annotation (∼15%), allowing for potentially more accurate prediction of function. *Tritrichomonas foetus* stands out, possessing 21 unique domains not found in the other species and exhibiting the highest number of unique domains overall, suggesting lineage-specific expansion or acquisition. By adding CAZyme annotation, 516 previously hypothetical or uncharacterised predicted genes now have potential functional assignments, improving genome annotation for these five trichomonads.

**Fig 1:**
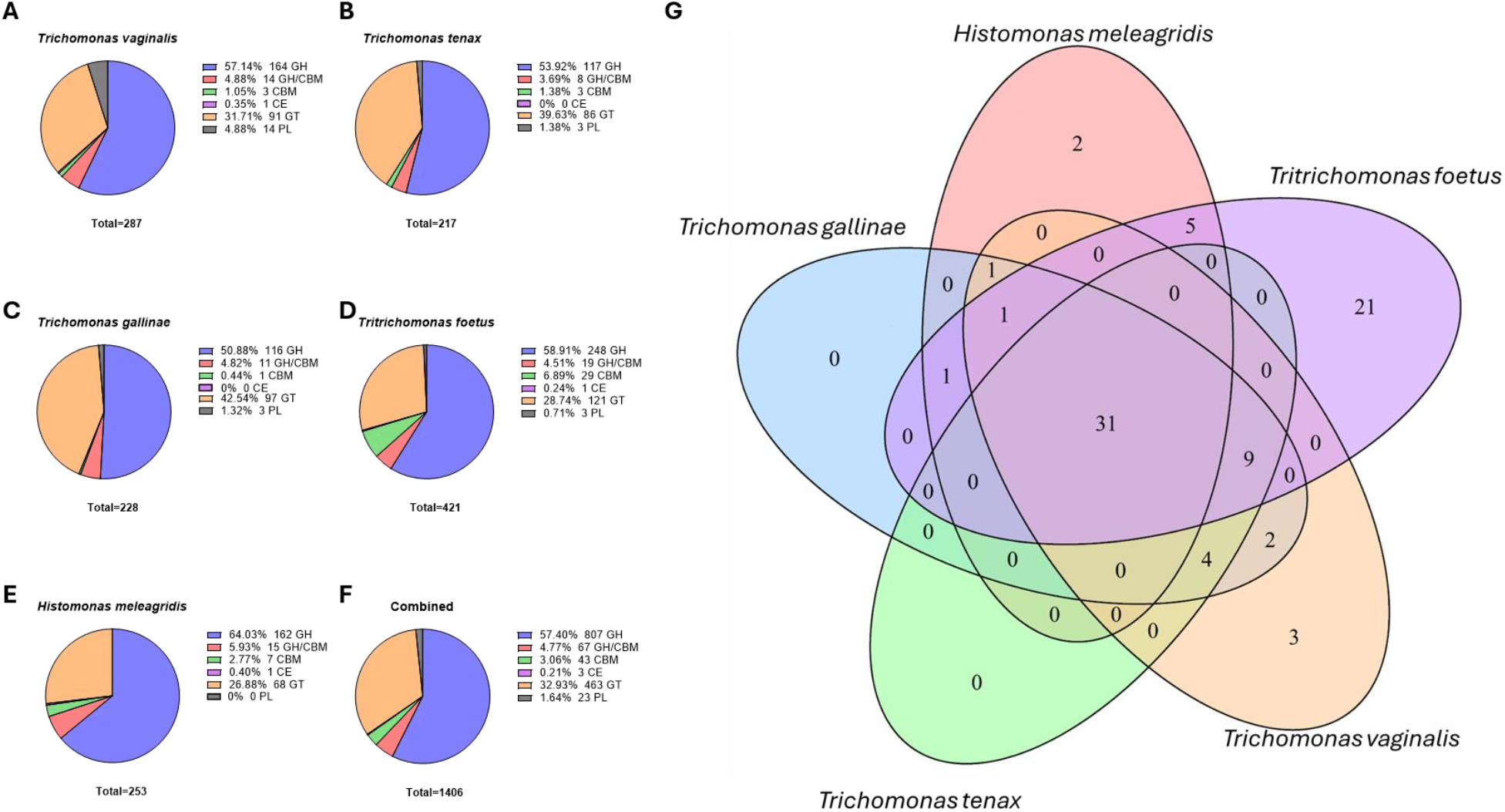
CAZy family composition and conservation across five trichomonad species. Pie charts show the total number and relative abundance of CAZyme families in each species (A–E) and across all five species combined (F). In the pie charts Glycoside hydrolases (GHs) are shown in blue; GHs containing carbohydrate-binding modules (GH/CBM) in red; carbohydrate-binding modules (CBMs) in green; carbohydrate esterases (CEs) in purple; glycosyltransferases (GTs) in orange; and polysaccharide lyases (PLs) in grey. A Venn diagram illustrates the conservation of CAZyme families among the five species (G), with each set labelled by its corresponding species.

Using Phobius, 310 (∼22%) of the 1,406 identified CAZymes have a predicted signal peptide (SP), consistent with secretion and potential extracellular activity (**Supplementary Table 1**). As it is difficult to predict SPs for divergent species like the trichomonads here it is pertinent to also attempt to predict SPs with different prediction software. Using SignalP-6.0 in eukaryotic mode, however, only 114 (∼8%) of the 1,406 CAZymes have a predicted SP. The majority of the CAZymes with a predicted SP by Phobius include a GH domain (216, ∼70%) (**Supplementary Table 1**). Phobius also predicts transmembrane domains (TMDs), suggesting that 495 (∼32%) of the identified CAZymes have one or more TMDs. Additionally, 298 (∼21%) of the 1,406 identified CAZymes are either predicted to have originated by ‘likely’ or ‘possible’ LGT (**Supplementary Table 1**) based on the Alienness score (Rancurel *et al*., 2017). The majority of those predicted to have likely originated via LGT are GHs (209/∼70%).

Several CAZyme families are highly duplicated, with more than ten counts detected across proteins from all five species analysed (**Table 2**). These include entries containing CBM20, GH13, GH31, GH47, and GT39, which are predominantly encoded by genes predicted by Alienness not to be of lateral origin. The CBM20 domain is typically involved in starch and glycogen binding and frequently occurs in tandem with enzymes from α-glucan active GH families (Christiansen *et al*., 2009; Zhu *et al*., 2014). One such domain, GH77 (a family that displays exclusively 4-α-glucanotransferase activity so far), was identified in 32 of the 36 proteins (∼89%) containing a CBM20 domain in the trichomonad genomes analysed here. Members of the GH13 family act on a range of substrates containing α-glucosidic linkages, including α-amylase activity (MacGregor *et al*., 2001), and, like GH77, are commonly associated with starch-binding CBM domains (Machovič & Janeček, 2006). Two GH13 subfamilies, GH13_8 and GH13_37, are conserved across all five trichomonads, with glycogen and starch as their respective candidate substrates. The conserved and duplicated presence of CBM20 and GH13 families highlights their likely functional importance in starch and glycogen binding and degradation in these organisms. GH31 family members, like GH13s, can exhibit α-glucosidase activity (Ikegaya *et al*., 2022). Subfamilies identified for trichomonad GH31s, GH31_1 and GH31_15, illustrate a likely function in hydrolysis of various α-(1→4), α-(1→6), or α-(1→3) linked polysaccharides and cleavage of nigerooligosaccharides (Ikegaya *et al*., 2022; Arumapperuma *et al*., 2023). All functionally characterised GH47 enzymes possess mannosidase activity, including the ability to cleave α-1,2-mannosidic linkages in N-glycans. These typically function in N-glycan maturation within the eukaryotic secretory pathway at the endoplasmic reticulum (Mast & Moremen, 2006; Xiang *et al*., 2016). Of the 94 GH47 proteins identified across the five trichomonad genomes, 12 (with at least one occurrence in each species) are predicted by Phobius to contain a signal peptide, and 49 are predicted to contain one or more TMDs, suggesting activity on extracellular substrates. These potential extracellular substrates have previously been hypothesised to include cell wall components of fungal members of the microbiota that share habitats with parasitic trichomonads, with cell walls containing an abundance of α-1,2-mannosidic linkages (Hall & Gow, 2013; Mpeyako *et al*., 2024). GH47-containing proteins comprise the largest orthogroup (OG0000000) identified in the orthogroup analysis, indicating strong conservation throughout the evolutionary history of the five trichomonads analysed.

**Table 2:** CAZy family counts per species for CAZy families conserved across all analysed genomes.

| CAZyme Family | *Trichomonas vaginalis* | *Trichomonas gallinae* | *Trichomonas tenax* | *Tritrichomonas foetus* | *Histomonas meleagridis* |
| --- | --- | --- | --- | --- | --- |
| CBM20 | 20 | 12 | 11 | 13 | 11 |
| CBM48 | 1 | 1 | 1 | 1 | 4 |
| GH1 | 2 | 2 | 2 | 3 | 5 |
| GH2 | 5 | 6 | 6 | 7 | 4 |
| GH3 | 4 | 6 | 7 | 18 | 6 |
| GH13 | 23 | 16 | 14 | 41 | 51 |
| GH14 | 4 | 4 | 4 | 6 | 19 |
| GH15 | 3 | 2 | 1 | 1 | 4 |
| GH19 | 8 | 5 | 7 | 1 | 8 |
| GH20 | 6 | 9 | 6 | 15 | 4 |
| GH30 | 16 | 1 | 1 | 3 | 2 |
| GH31 | 17 | 12 | 17 | 20 | 12 |
| GH36 | 4 | 4 | 2 | 16 | 2 |
| GH38 | 5 | 4 | 4 | 8 | 2 |
| GH47 | 28 | 23 | 24 | 9 | 10 |
| GH77 | 14 | 6 | 7 | 17 | 6 |
| GH99 | 6 | 7 | 8 | 5 | 2 |
| GH133 | 2 | 2 | 2 | 3 | 4 |
| GH163 | 10 | 2 | 2 | 1 | 4 |
| GT1 | 2 | 2 | 2 | 1 | 2 |
| GT2 | 5 | 4 | 6 | 23 | 4 |
| GT4 | 2 | 2 | 2 | 2 | 6 |
| GT5 | 1 | 1 | 1 | 6 | 2 |
| GT8 | 8 | 13 | 6 | 4 | 4 |
| GT24 | 9 | 9 | 8 | 10 | 8 |
| GT33 | 3 | 2 | 3 | 3 | 2 |
| GT35 | 2 | 2 | 2 | 9 | 6 |
| GT39 | 32 | 28 | 30 | 23 | 10 |
| GT47 | 8 | 7 | 5 | 8 | 8 |
| GT61 | 4 | 6 | 4 | 6 | 8 |
| GT66 | 5 | 2 | 3 | 9 | 4 |

### Tritrichomonas foetus-specific CAZymes

As described above, *Tritrichomonas foetus*, encodes an additional 21 unique CAZy families compared with the other four parasitic trichomonads analysed here (**Table 3**). Several of these unique *Tritrichomonas foetus* CAZy families are highly duplicated (≥6 family occurrences), including CBM51, GH35, GH95, and GT10.

**Table 3:** CAZy family counts for *Tritrichomonas foetus*-specific CAZy families, including predicted acquisition by lateral gene transfer (LGT) and representative functions.

| CAZyme Family | *Tritrichomonas foetus* | Predicted LGT (Y/N) | Example Function |
| --- | --- | --- | --- |
| CBM51 | 6 | N | Galactose Binding |
| CBM52 | 1 | Y | β-1,3-glucan Binding |
| GH35 | 13 | N | β-galactosidase |
| GH43 | 4 | N | α-L-arabinofuranosidase |
| GH65 | 1 | N | maltose (Glc-α-1,4-Glc) phosphorylase |
| GH76 | 3 | Y | α-mannanase |
| GH78 | 3 | N | α-L-rhamnoside |
| GH81 | 1 | Y | endo-β(1,3)-glucanase |
| GH89 | 3 | N | α-N-acetylglucosaminidase |
| GH95 | 8 | N | 1,2-α-L-fucosidase |
| GH125 | 3 | N | α-mannosidase |
| GH128 | 1 | Y | endo-β-1,3-glucanase |
| GH130 | 1 | Y | 4-O-β-D-mannosyl-D-glucose phosphorylase |
| GH132 | 1 | Y | exo-β-1,3-glucosidase |
| GH141 | 1 | N | α-L-fucosidase |
| GT10 | 6 | N | 4-galactosyl-N-acetylglucoside 3-α-L-fucosyltransferase |
| GT14 | 1 | Y | β-1,3-galactan β-1,6-glucuronosyltransferase |
| GT17 | 1 | N | β-1,4-mannosyl-glycoprotein β-1,4-N-acetylglucosaminyltransferase |
| GT75 | 1 | N | self-glucosylating β-glucosyltransferase |
| CE9 | 1 | N | deacetylation of N-acetylglucosamine-6-phosphate to glucosamine-6-phosphate |
| PL9 | 2 | Y | pectate lyase |

Two of the unique to *Trichomonas foetus* CAZy families, GH76 and GH125, are both α-mannan active with trichomonad GH76 family members, which are particularly common in bacteria, associated with the degradation of fungal α-mannans in the fungal cell wall (Cuskin *et al*., 2015). The GH76s identified in *Tritrichomonas foetus* are classified as LGT candidates, with closest homologues in *Parabacteroides spp.*, consistent with acquisition from Bacteroidales bacteria (Maruyama & Nakajima, 2000; Cuskin *et al*., 2015). The expanded repertoire of mannan-related CAZymes in *Tritrichomonas foetus* may therefore facilitate the targeting of mannans derived from co-occurring microbiota.

One of the relatively few PLs, PL9, identified in this study, appears to be specific to *Tritrichomonas foetus*. These PL9s are classified as LGT candidates, with best matches in *Bacteroides* species **(Supplementary Table 1)**. In human colonic *Bacteroides* species, such as *Bacteroides thetaiotaomicron*, PL9 enzymes contribute to the degradation of dietary pectic glycans in conjunction with other GHs and PLs, with the corresponding genes organised into polysaccharide utilisation loci (PULs) (Luis *et al*., 2018). Plant pectins play an important role in maintaining a healthy, eubiotic gut microbiota, and alterations to their structural properties can influence the composition of mutualistic bacterial populations (Larsen *et al*., 2019). Moreover, pectin as a dietary supplement is an effective prebiotic, where supplementation enhances the relative abundance of protective *Lactobacillus* species in the vaginal microbiota in mammals (He *et al*., 2024). From an infection perspective, the ability to modify or degrade pectin could represent a mechanism by which *Tritrichomonas foetus* could perturb or sustain dysbiosis within its niche (bovine urogenital tract); however, further experimental work is required to test this hypothesis. But pectins would be expected to be present in the gut rather than the urogenital tract. Hence, these data could be considered to be consistent with the gut location of *T. foetus* in cats, dogs and pigs (Tachezy *et al*., 2002; Dąbrowska *et al*., 2019) as well as more recently shown for cattle (Martínez *et al*., 2023).

### Putative Host Mucin O-Glycan Targeting CAZymes

The capacity for mucin targeting by CAZymes in *Trichomonas vaginalis* and *Trichomonas tenax* has previously been inferred, with several putative mucin O-glycan–degrading CAZymes identified and shown to be conserved between these two species (Labourel *et al*., 2023; Raba & Luis, 2023; Mpeyako *et al*., 2024). Several of these CAZyme families are also conserved across all five trichomonad species analysed here, specifically GH20 and GH2 (**Figure 2**). Each of these families includes enzymes that can degrade core structures of mucin O-glycans in other organisms (Bakshani *et al*., 2025), suggesting similar potential in these trichomonads. Notably, *Trichomonas* and *Tritrichomonas* species exhibit a broader repertoire of conserved, putatively mucin-targeting CAZymes. The conservation of a diverse set of mucin-targeting CAZymes across these four species is consistent with their capacity for host and site switching between distinct mucosal environments (Dąbrowska *et al*., 2020; Bailey *et al*., 2023; Sullivan *et al*., 2025), where mucin O-glycan composition varies. There is evidence of host and site switching for *Histomonas meleagridis* as well (Beer *et al*., 2022), so host/site switching alone may not be sufficient selection pressure to maintain a broad mucin-targeting GH repertoire, potentially reflecting differences in nutrient acquisition strategies, reliance on co-occurring microbiota, or lineage-specific gene loss. These GH families display complex evolutionary histories, with the same GH classes inferred to have been acquired via lateral gene transfer in some trichomonad lineages while being vertically inherited in others. For example, GH20 CAZymes in *Trichomonas* species were likely acquired through lateral transfer from prokaryotes based on the Alienness analysis, whereas GH20s in *Tritrichomonas foetus* and *Histomonas meleagridis* appear to have been retained through vertical inheritance. In total, 209 putative mucin O-glycan–degrading CAZymes were identified across the five genomes analysed (**Figure 2**). Of these, 77 proteins (∼37%) are predicted by Phobius to contain a signal peptide, consistent with activity on extracellular substrates. Using phylogenetic methods suggests that the GH2 enzymes from all the trichomonads analysed here could have been acquired by lateral transfer from prokaryotes however originating from multiple different acquisition events (**Figure 3**). These findings underscore the importance of experimentally characterising the mucin-degrading capacities of these trichomonads to determine their potential roles in dysbiosis and disease at the mucosal sites they infect.

**Fig 2:**
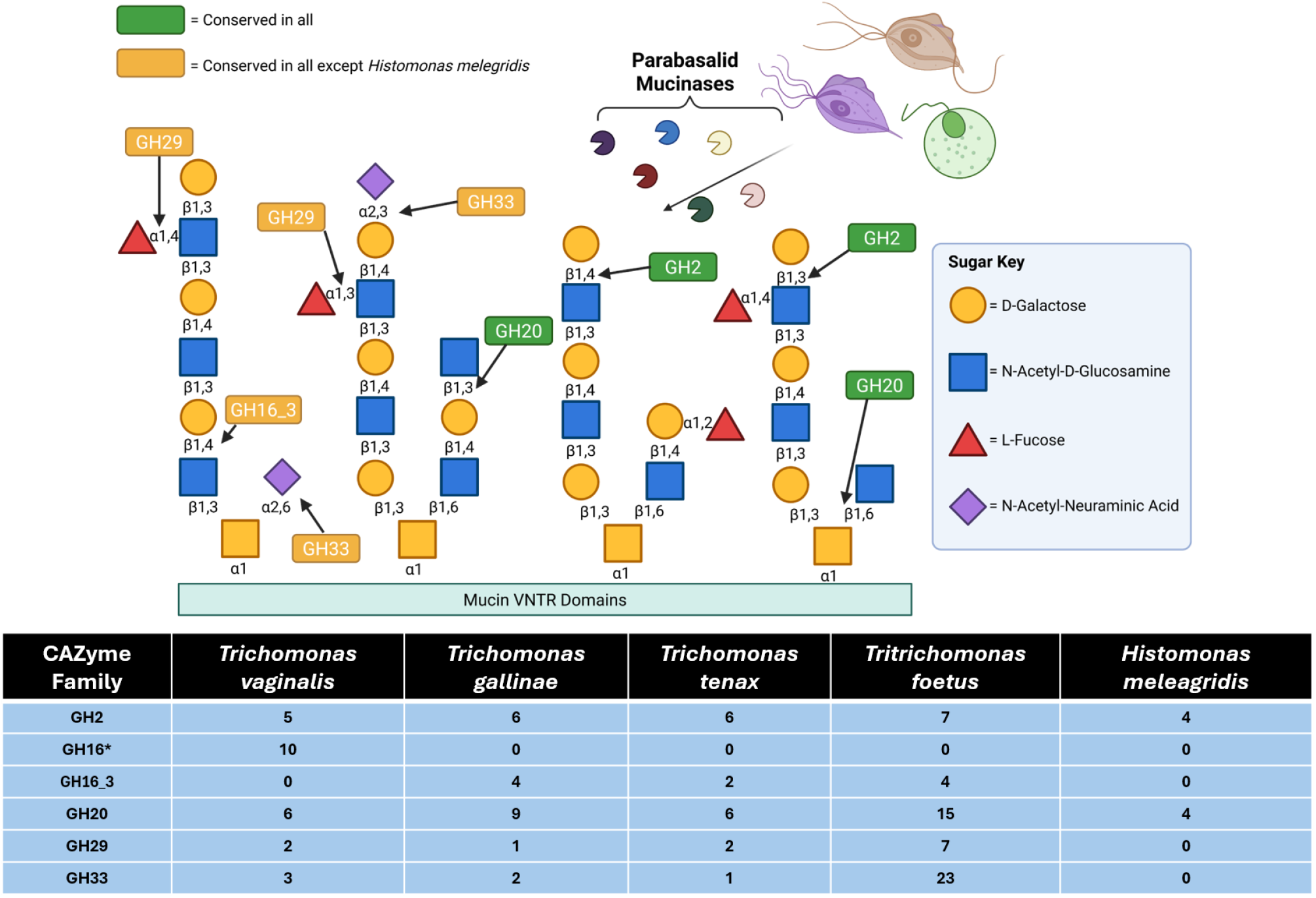
Example mucin O-glycans and their GHs, and a diagrammatic representation of trichomonad mucinases. Arrows indicate the glycosidic bonds known to be hydrolysed by individual GHs. GHs shown in green boxes are present in all five trichomonad genomes, whereas those shown in yellow boxes are absent from Histomonas meleagridis but present in the other four species. Sugar components of mucin chains are depicted using the Symbol Nomenclature for Glycans (Varki *et al*., 2015). The table below shows the number of candidate O-glycan-degrading GHs in each species. *The GH16s found in Trichomonas vaginalis do not have sub family annotation however bare significant similarity to all other Trichomonad GH16s which have the subfamily classification GH16_3. Adapted from Labourel et al., 2023, Raba and Luis, 2023 and Mehlhorn, 2015.

**Fig 3:**
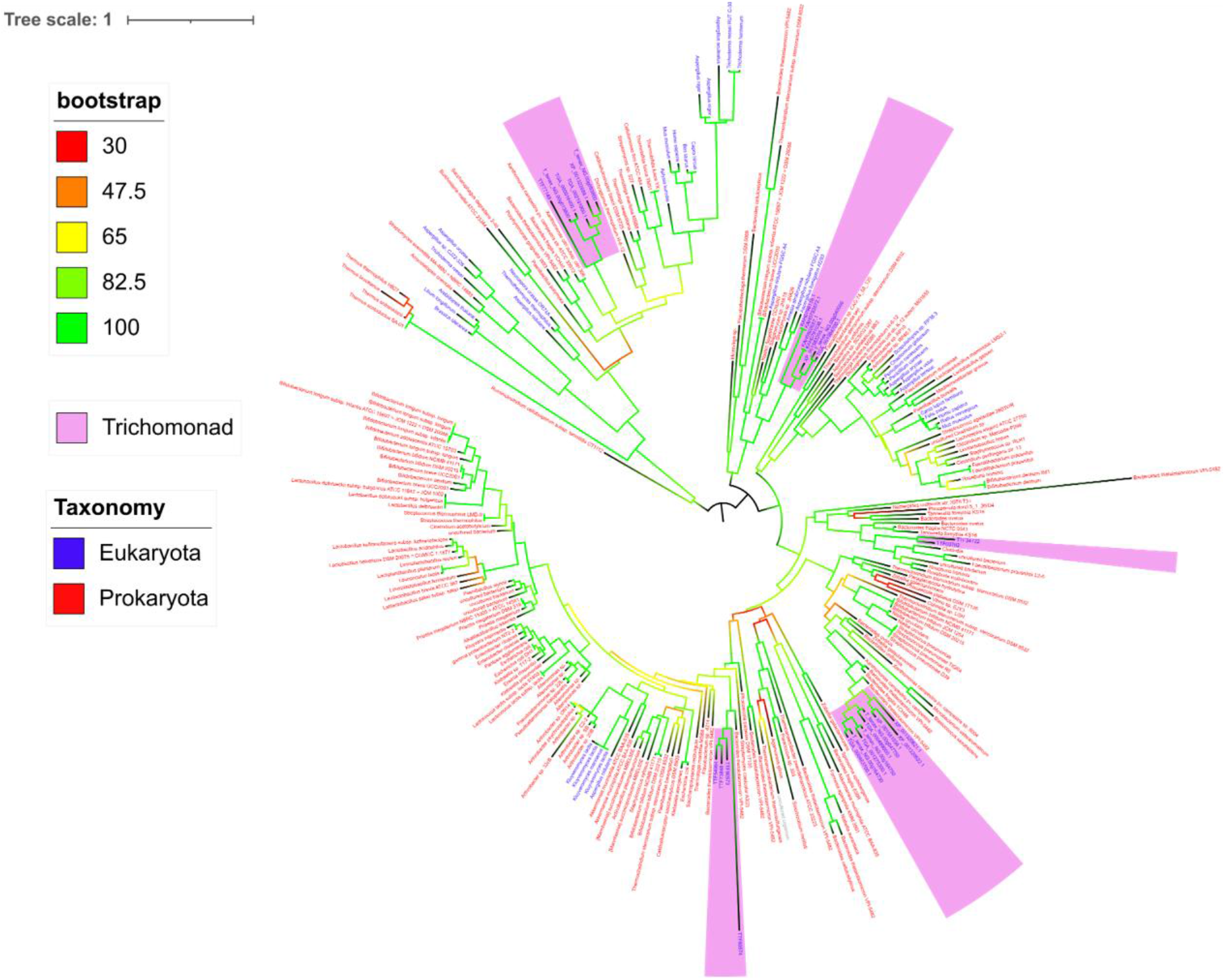
Midpoint-rooted phylogeny of characterised GH2 sequences from the CAZy database and metamonad GH2 sequences. Trichomonad GH2 sequences are highlighted in purple. Terminal-node labels include the species name and sequence identifier; labels are coloured blue for eukaryotic sequences and red for prokaryotic sequences. Branch colours represent bootstrap support, with greener branches indicating higher support and redder branches indicating lower support.

### Laterally Acquired Antibacterial Toolkit Conservation

Using a combination of co-culture transcriptomics and targeted molecular biology approaches, a laterally acquired antibacterial enzyme toolkit - predominantly composed of CAZymes targeting the bacterial cell wall - was previously identified in *Trichomonas* species (Hart *et al*., 2025). Phylogenetic analyses suggested that many of these CAZymes are conserved across trichomonads. The re-annotation performed in this study supports these findings, demonstrating that all five considered species GH19 muramidases and putative N-acetylglucosaminase GH3 enzymes, with subsets of these as likely LGT candidates (**Figure 4**). However, some GH3 enzymes in *Trichomonas* species are likely distinct from those in *Tritrichomonas foetus* and *Histomonas meleagridis*: only a subset of *Trichomonas* GH3s are classified as lateral gene transfer candidates by alienness, and these cluster into different orthogroups from GH3s in *Tritrichomonas foetus* and *Histomonas meleagridis*, consistent with previous phylogenetic analyses (Hart *et al*., 2025). Phylogenetic analysis in this work, however using sequences from all characterised GH3 enzymes in the CAZy database, shows that the trichomonad GH3s identified in our annotation cluster into several distinct clades all clustering with prokaryotes. This suggests multiple independent acquisition events and indicates functional diversity among trichomonad GH3s (**Figure 5**), potentially highlighting a weakness in the alienness methodology compared with more precise phylogenetic approaches. In addition, *Tritrichomonas foetus* and *Histomonas meleagridis* lack the laterally acquired muramidase GH25 CAZymes present in *Trichomonas* species. Collectively, these patterns suggest that the capacity to degrade bacterial cell wall polysaccharides is broadly conserved across trichomonads and is likely functionally important in their interactions with mucosal microbiota. However, *Trichomonas* species exhibit greater redundancy in muramidase activity, potentially reflecting heightened selective pressure for antibacterial functionality in these lineages. Nevertheless, the presence of these CAZymes suggests that all five species analysed in this study are capable of dynamically interacting with bacterial members of their respective mucosal microbiota, with potential consequences for the initiation or modulation of dysbiosis and disease, although the precise in vivo impact of these enzymes remains to be determined.

**Fig 4.**
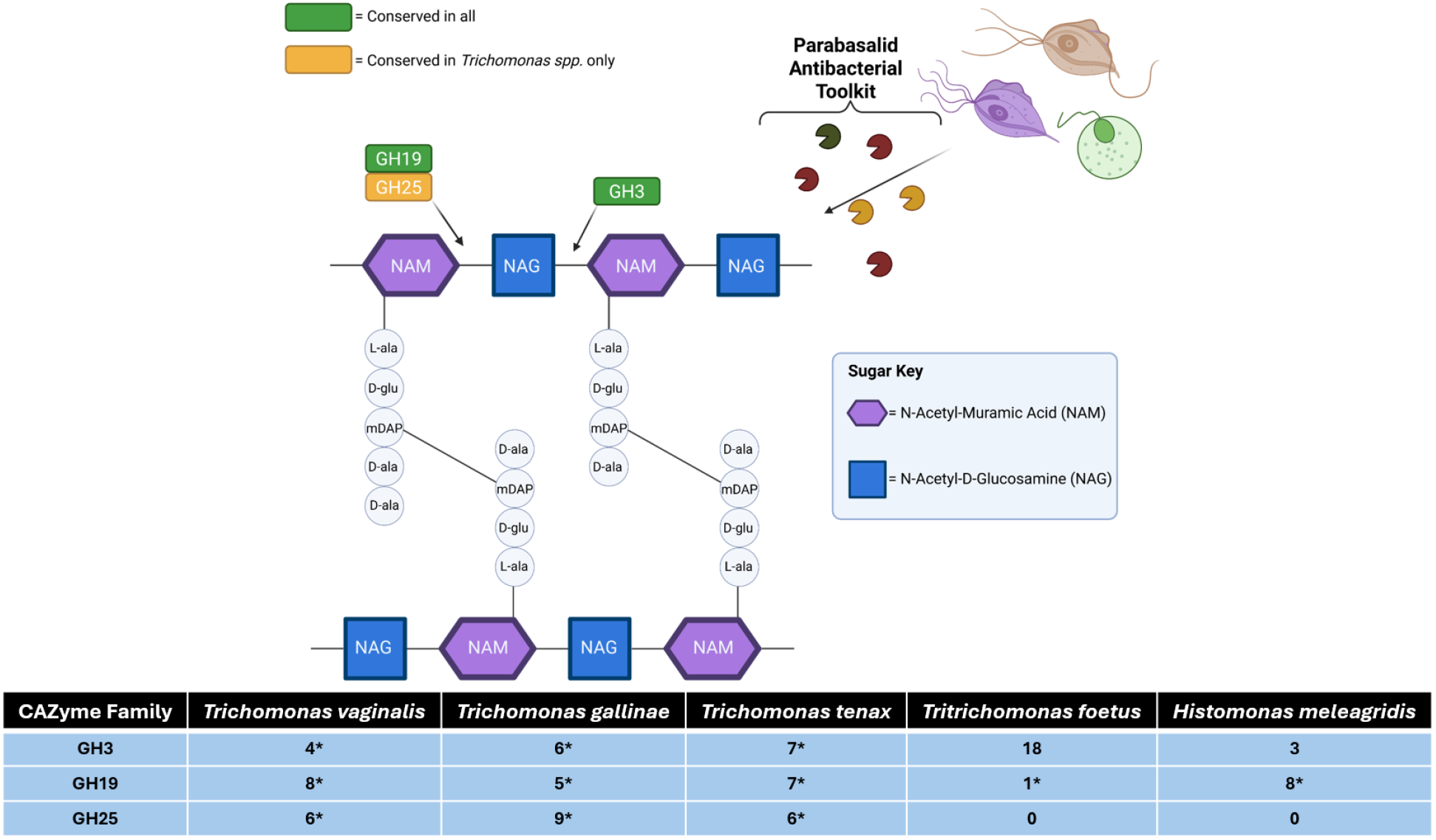
Example peptidoglycan structure, its associated glycoside hydrolases (GHs), and a schematic representation of trichomonad peptidoglycan hydrolases. Arrows indicate glycosidic bonds known to be hydrolysed by individual GHs. GHs shown in green boxes are present in all five trichomonad genomes, whereas those shown in yellow boxes are absent from *Histomonas meleagridis* and *Tritrichomonas foetus* but present in the other three species. The table below shows the number of candidate peptidoglycan-hydrolysing GHs in each species. GHs marked with an asterisk in the table are predicted to have been acquired by lateral gene transfer. Although GH3 enzymes are active against the indicated β-GlcNAc bond, they cleave only pre-digested peptidoglycan products generated by muramidase activity.

**Fig 5:**
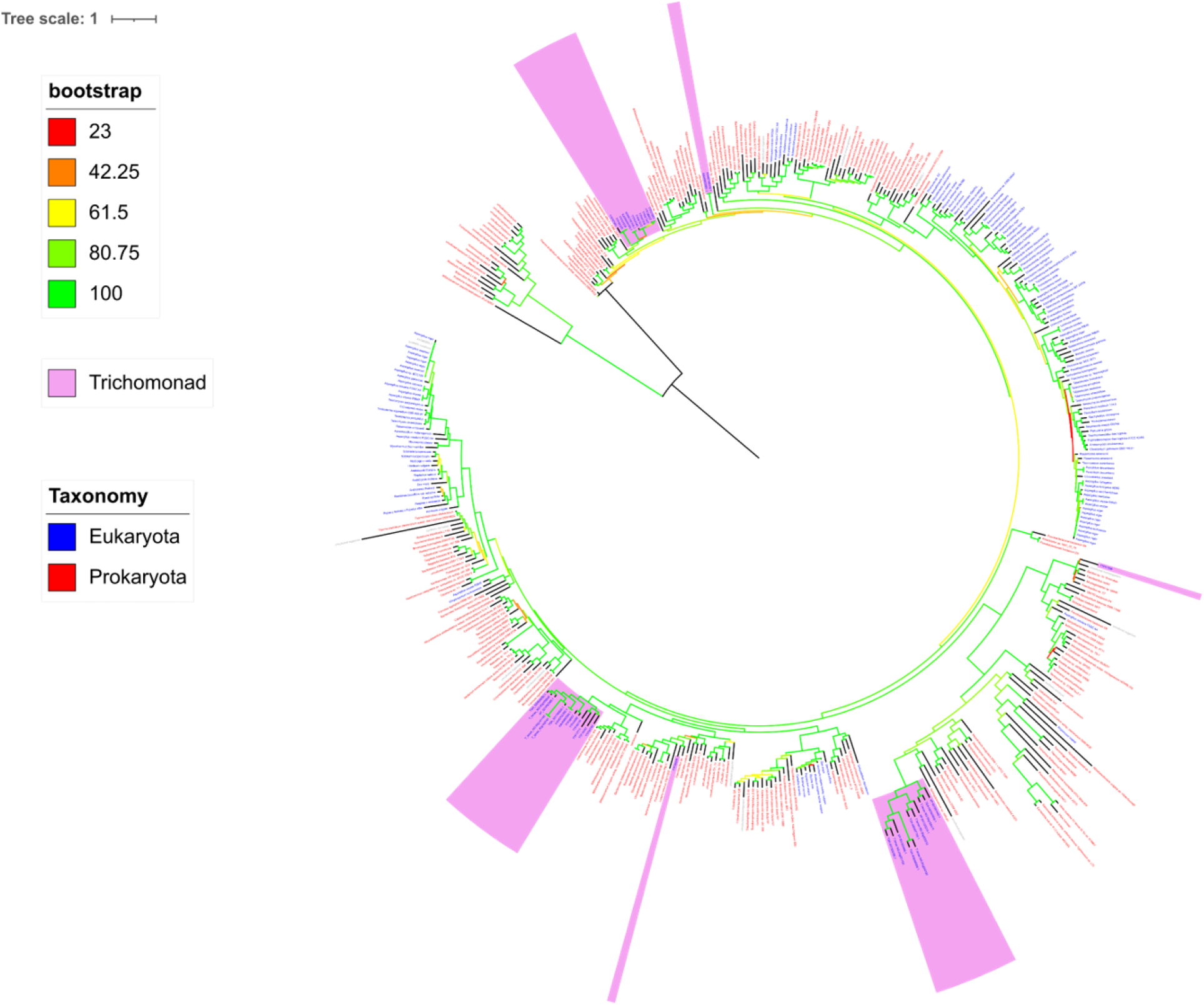
Midpoint-rooted phylogeny of characterised GH3 sequences from the CAZy database and trichomonad GH3 sequences. Trichomonad GH3 sequences are highlighted in purple. Terminal-node labels include the species name and sequence identifier; labels are coloured blue for eukaryotic sequences and red for prokaryotic sequences. Branch colours represent bootstrap support, with greener branches indicating higher support and redder branches indicating lower support.

## Discussion

Parasitic trichomonads are associated with or are a primary causative agents of disease in mammals and birds (Lawson *et al*., 2011; Kissinger & Adamski, 2013; Amin *et al*., 2014; Mehlhorn, 2014; Dąbrowska *et al*., 2019). Their genomes are unusually large for microbial eukaryotes/protists and characterised by extensive gene duplication, large-scale prokaryote-to-eukaryote lateral gene transfer, and a high proportion of poorly annotated or hypothetical genes, complicating the identification of putative virulence factors. The CAZyme repertoires of organisms inhabiting mucosal surfaces have been implicated in virulence and disease, including through the modification of host mucins and bacterial cell walls in trichomonad species (Stafford *et al*., 2017; Labourel *et al*., 2023; Raba & Luis, 2023; Mpeyako *et al*., 2024; Hart *et al*., 2025). Accordingly, this study aimed to augment the annotation of five disease-associated trichomonad genomes by systematically identifying CAZy entries, thereby facilitating the discovery of putative virulence factors and other disease-relevant genes and to provide an integrated resource that links CAZyme presence, predicted origin, and subcellular localisation to hypotheses about trichomonad interactions with host and host-associated microbiota. In total, CAZyme annotations were assigned to 1,406 unique protein-coding genes, including 516 previously hypothetical proteins lacking informative annotation (**Figure 1**). In addition, LGT predictions and accessory annotations, such as signal peptide and TMD predictions and KEGG pathway assignments, were generated, although these remain inference-based and require experimental validation. Together, these resources will enable researchers to identify genes of interest amongst trichomonad CAZymes and CBMs.

To identify CAZymes that are evolutionarily conserved among trichomonads - and by extension, those likely fundamental to their mucosal lifestyles - we examined CAZy classes present across all five species in greater detail. In total, 31 CAZyme domain families were detected at least once in each genome (**Table 2**). Several of these families were extensively duplicated, with more than ten family copies per genome, consistent with functional importance and a selective advantage of redundancy. These highly duplicated families include CBM20, GH13, GH31, GH47, and GT39. Functionally, these CAZymes and CBMs are predicted, based on characterised homologues in other organisms, to participate in the binding and degradation of α-glucans, including starch and glycogen, potentially derived either from the host or from cohabiting members of the mucosal microbiota (Mpeyako *et al*., 2024). *Tritrichomonas foetus* exhibited an expanded CAZyme repertoire relative to the other four trichomonads, encoding 21 additional CAZyme domain families unique to this species (**Table 3**). Notably, several of these classes are associated with α-mannan processing, indicating that *Tritrichomonas foetus* may possess increased functional redundancy for mannan utilisation compared with other trichomonads.

To explore putative virulence factors, this study examined in detail a conserved repertoire of CAZy domains predicted to target host mucin O-glycans (**Figure 2**). All five trichomonads encode CAZymes potentially capable of hydrolysing core mucin O-glycan structures, specifically GH2 and GH20 family members. Additional putative mucin O-glycan–degrading capacity is predicted for all species except *Histomonas meleagridis*, which lacks GH16_3, and GH33, despite retaining the core GH2 and GH20 families. It should be noted, however, that in the absence of functional validation, these predicted GHs may act on alternative substrates and possess activities unrelated to mucin O-glycan degradation, as many CAZy families encompass enzymes with diverse specificities. Notably, the majority of these putative mucin-targeting CAZymes in *Trichomonas* species appear, based on Alienness scores, to have been acquired via lateral gene transfers, whereas the same CAZy domains in *Histomonas meleagridis* and *Tritrichomonas foetus* are predominantly vertically inherited **(Supplementary Table 1)**. This pattern suggests complex evolutionary histories for these families, with similar putative mucin-targeting capacities arising via different combinations of lateral acquisition and vertical retention across trichomonad lineages, provided that CAZyme family functions are conserved. Given that mucin degradation by mucosal pathobionts is frequently associated with disease and the identification of conserved, duplicated, and occasionally laterally acquired putative mucin hydrolases together highlights an important avenue for future functional investigation in these organisms.

Consistent with previous phylogenetic analyses, all five trichomonads show conservation of CAZymes capable of degrading the polysaccharide backbone of bacterial cell wall peptidoglycan (Hart *et al*., 2025). These include muramidases (GH19 and putatively GH25) and putative N-acetylglucosaminases (GH3), which together can potentially cleave both glycosidic linkages within peptidoglycan (Vollmer *et al*., 2008). All of these enzymes (GH19, GH25, GH3) are classified as lateral gene transfer candidates either by alienness or phylogenetics or both, consistent with previous phylogenetic analyses (Hart *et al*., 2025). However, *Histomonas meleagridis* and *Tritrichomonas foetus* exhibit reduced functional redundancy, as they lack GH25 CAZymes, potentially conferring comparatively lower muramidase activity. In addition to these CAZymes, peptidase enzymes capable of cleaving peptide cross-links within peptidoglycan have been described in trichomonads and functionally characterised in *Trichomonas vaginalis*. These enzymes likely act in concert with peptidoglycan-hydrolysing CAZymes as part of an antibacterial toolkit targeting bacterial cell wall components (Pinheiro *et al*., 2018; Barnett *et al*., 2023; Hart *et al*., 2025). The likely coordinated activity of these enzymes could therefore influence the initiation or maintenance of dysbiosis within mucosal microbiotas by potentially targeting mutualistic bacterial community members and contributing to inflammation through generating strong pro-inflammatory peptidoglycan fragments (Hart *et al*., 2025).

As with all in silico genome annotation efforts, experimental functional validation remains essential to confirm predicted gene functions and activities. The CAZy domain annotations presented here provide such a foundation, prioritising over 1,400 carbohydrate-active genes - including 516 previously hypothetical proteins - for targeted biochemical and infection-model studies. These CAZymes, many of which are conserved across all five considered species, are extensively duplicated and shaped by lateral gene transfers, and represent promising candidates whose functional characterisation will clarify how parasitic trichomonads exploit host and microbial glycans to drive virulence, dysbiosis, and disease at mucosal sites.

## Methods

### CAZy Annotation

Briefly, CAZyme families were predicted for all previously identified protein-coding genes from five trichomonad genomes (*Histomonas meleagridis* (2922-C6/04-10x), *Trichomonas gallinae* (A1), *Trichomonas tenax* (Hs-4:NIH/ATCC 30207), *Trichomonas vaginalis* (G3) and *Tritrichomonas foetus* (Belfast)) based on significant similarities with entries from the curated CAZy database (Drula *et al*., 2022). For borderline cases and suspected fragments, manual curation was also applied to the CAZy family annotations, as previously done for numerous entries in the CAZy database (Lombard *et al*., 2014).

### Alienness prediction

For identified trichomonad CAZymes, the likelihood that a given gene originated via lateral gene transfer was assessed using the Alienness method (Rancurel *et al*., 2017). Homologous sequences were identified using DIAMOND blastp (v2.1.18) against the UniRef90 protein database, with a maximum of 500 target sequences, e-value < 1e-5, and the ‘sensitive’ mode. (Buchfink *et al*., 2021). NCBI taxonomic identifiers for DIAMOND hits were retrieved using the UniProt ID mapping tool and appended to the DIAMOND tabular output. The resulting dataset was uploaded to the Alienness server, where predictions were performed with Parabasalia (taxid: 5719) specified as the focal taxonomic group. Taxa corresponding to trichomonad species which have had added CAZyme annotation in this work were excluded from the analysis (*Histomonas meleagridis*, taxid: 135588; *Trichomonas gallinae*, taxid: 56777; *Trichomonas tenax*, taxid: 43075; *Trichomonas vaginalis*, taxid: 5722; and *Tritrichomonas foetus*, taxid: 1144522).

### KEGG annotation

KEGG annotation was added to all trichomonad CAZymes using the BlastKOALA tool and retaining the best KO hits (Kanehisa *et al*., 2016).

### Orthologous gene identification

The orthogroups of trichomonad CAZymes were inferred using OrthoFinder v2.5.5 (Emms & Kelly, 2015; Emms & Kelly, 2019) from only the Trichmonad CAZymes as an input.

### Signal Peptide Identification

Signal peptides (and transmembrane domains) were predicted for all trichomonad CAZymes using SignalP 6.0, using the eukaryotic model on default parameters (Nielsen *et al*., 2024), and Phobius (Käll *et al*., 2004).

### Phylogenetic Analysis

Sequences for multiple sequence alignments were retrieved from the CAZy database, including only entries annotated as characterised CAZymes. Alignments were generated, curated, and masked using SeaView (Gouy *et al*., 2009). Phylogenetic trees were inferred from the masked alignments, with emphasis on conserved regions, using the IQ-TREE web server (Trifinopoulos *et al*., 2016), and subsequently visualised and annotated with iTOL (Letunic & Bork, 2024).

## Data Availability

The data underlying this article are available in the article and in its online supplementary material.

## Acknowledgements

A.J.H. was supported by the UK Biotechnology and Biological Sciences Research Council Doctoral Training Partnership program for Newcastle, Liverpool, and Durham (Grant number: BB/T008695/1, R.P.H. supervisor) and UK Biotechnology and Biological Sciences Research Council Future Leaders Fellowships (Fellowship number: MR/V027204/1, M.B. fellowship recipient).

